# HiCP2GAN: A Plug and Play Foundation Model-based GAN for Hi-C Enhancement

**DOI:** 10.64898/2026.05.18.725960

**Authors:** Samuel Olowofila, Oluwatosin Oluwadare

## Abstract

The three-dimensional organization of chromatin shapes gene regulation and cellular function. Hi-C has emerged as the primary technique for mapping chromatin interactions genome-wide, yet high-resolution data remain costly and scarce, leaving many studies with sparse contact maps that limit downstream analysis. Deep learning methods, especially generative adversarial networks (GANs), have shown promise for enhancing low-resolution Hi-C data. Most existing GAN-based approaches, however, rely on custom discriminators trained from scratch, which can yield unstable training and limited generalization. Hi-C foundation models pretrained on large-scale data capture rich, transferable representations of chromatin structure; their use as discriminators within adversarial enhancement frameworks has not been explored. In this work, we introduce HiCP2GAN, a plug-and-play GAN that employs a pretrained Vision Transformer-based Hi-C foundation model as its discriminator. The discriminator was pretrained on 118 million Hi-C patches across diverse species and cell types, providing biologically meaningful gradients for adversarial supervision. The HiCP2GAN framework is generator-agnostic: any compatible Hi-C resolution enhancement architecture can serve as the generator, enabling fair comparison across methods. The encoder phase of the foundation model was adapted as a discriminator backbone and experimented with finetuning different numbers of layers from the input while freezing the deeper transformer layers. Finetuning the first few layers while freezing the rest preserved pretrained knowledge while allowing task-specific adaptation. Experiments on human cell lines show that HiCP2GAN consistently improves resolution over standalone generators and conventional GAN-based models, while serving as a plug-and-play framework for most non-GAN generator models. HiCP2GAN is publicly available at https://github.com/OluwadareLab/HiCP2GAN.

## 1 Introduction

The spatial organization of chromatin in the cell nucleus plays a cardinal role in modeling the gene regulatory landscape. Hi-C, the prime high-throughput chromosome conformation capture technique for genome-wide assessment of chromatin interaction frequencies(IF) provides critical insights into higher-order chromatin structures, loops, and topological associating domains(TADs) (Lieberman-Aiden *et al*., 2009). However, the required high-resolution (HR) data across cell types and experimental conditions are inherently constrained by sequencing depth, experimental cost, and data noise.

A growing body of research has focused on mitigating these limitations using computational super-resolution techniques. Early approaches leveraged statistical modeling ((Imakaev *et al*., 2012)), while later methods explored deep learning frameworks including Convolutional Neural Networks (CNNs) such as HiCNN1 (T. Liu and Z. Wang, 2019a), HiCPlus (Zhang *et al*., 2018), HiCNN2 (T. Liu and Z. Wang, 2019b), HiCARN1 (Hicks and Oluwadare, 2022), SRHiC (Z. Li and Dai, 2020), and HiCPlus (Zhang *et al*., 2018); Generative Adversarial Networks (GANs) (Goodfellow *et al*., 2020) inclusive of HiCSR (Dimmick, 2020), HiCARN2 (Hicks and Oluwadare, 2022), HiCGAN (Q. Liu *et al*., 2019) DeepHiC (Hong *et al*., 2020), and EnHIC (Hu and W. Ma, 2021), and Attention-based mechanisms such as DiCARN (Olowofila and Oluwadare, 2025) and (P. Li *et al*., 2025).

In spite of these advances, most Hi-C resolution enhancement methods still suffer from constraints, including monolithic architectures, tightly coupled generator designs, resolution hallucination and assumptions, normalization strategies, and loss function mismatching. As a result, direct comparison across methods is confounded by incompatible workflows, differing output ranges, and inconsistent evaluation protocols. Also, the majority of GAN-based approaches exhibit shallow convolutional discriminators that essentially capture local feature data, resulting in limited abilities to capture long-range chromatin dependencies that are crucial to 3D genome organization and analyses.

Several GAN-based methods have been proposed for Hi-C resolution enhancement (Table I). hicGAN (Q. Liu *et al*., 2019) adapts SRGAN to Hi-C matrices, using a skip-connection generator and purely adversarial loss, making it susceptible to missing fine-scale details. DeepHiC (Hong *et al*., 2020) combines MSE, total variation, perceptual (VGG-derived), and adversarial losses, but perceptual loss can introduce unwanted image-like textures (Tej *et al*., 2020). HiCSR (Dimmick, 2020) uses a skip-connection generator, a CNN discriminator, and feature loss from a denoising autoencoder. HiCARN2 (Hicks and Oluwadare, 2022) and EnHiC (Hu and W. Ma, 2021) introduce domain-specific designs: HiCARN2 uses residual blocks tailored to Hi-C, while EnHiC exploits rank-1 matrix decomposition to better match Hi-C structure. <u>A common limitation of these methods is that their discriminators are trained from scratch and are typically shallow CNNs, limiting their ability to encode long-range chromatin topology</u>.

**Table I:** Overview of GAN-based Hi-C enhancement methods. Prior approaches rely on convolutional discriminators trained from scratch, whereas HiCP2GAN replaces the discriminator with a pretrained ViT-based Hi-C foundation model and remains compatible with any generator architecture.

| Method | Discriminator | Foundation model |
| --- | --- | --- |
| hicGAN | CNN | No |
| DeepHiC | CNN | No |
| HiCSR | CNN | No |
| HiCARN2 | CNN | No |
| EnHiC | CNN | No |
| HiCP2GAN (Ours) | ViT (pretrained) | Yes |

In recent computational evolution, foundation models (C. Ma *et al*., 2024; H. Li *et al*., 2025), typically pretrained on very large domain-centered datasets and have demonstrated remarkable success across various domain-specific tasks ranging from computer vision to natural language processing, and more importantly, biological modalities. The Hi-C analyses domain has recently recorded the encoding of rich chromatin structure representation when using transformer-based architectures pretrained on diverse contact maps, demonstrating the global interaction patterns adaptable for varying downstream exploration (X. Wang *et al*., 2024).

In this study, we introduce HiCP2GAN, a novel generator-agnostic generative adversarial method for Hi-C reconstruction that integrates a pretrained Hi-C foundation model as its backbone discriminator. HiCP2GAN tackles the Hi-C resolution enhancement problem with foundation-supervised adversarial learning, where various generator architectures are evaluated and optimized under a unified, semantically nuanced discriminator.

One key idea underlying HiCP2GAN is the decoupling of the generator construction from its adversarial supervision. By enforcing a standardized input-output interface, consistent numerical representation, and shared discriminator architecture, HiCP2GAN enables fair, controlled comparison across heterogeneous super-resolution methods. The adopted foundation-based discriminator (X. Wang *et al*., 2024) uses self-attention (Vaswani *et al*., 2017) to capture long-range chromatin interactions, thus providing adversarial feedback that adequately reflects biologically meaningful structural properties instead of purely local signal fidelity.

We demonstrate that foundation-supervised adversarial training consistently improves Hi-C reconstruction while revealing inherent architectural trade-offs as well as strengths amongst various generator models. HiCP2GAN could now serve as both a benchmarking framework and a generalizable training scaffold for future Hi-C super-resolution methods.

## 2 Materials and Methods

### 2.1 HiCP2GAN Framework Overview

HiCP2GAN is an adversarial super-resolution framework designed to enhance the spatial fidelity of low-resolution (LR) Hi-C contact matrices while remaining agnostic to the underlying generator architecture. Unlike prior Hi-C enhancement models that tightly couple architectural design, resolution assumptions, and loss formulations into a single model, HiCP2GAN decouples generator construction from discriminator supervision through the integration of a foundation-level representation learner.

The framework consists of three principal components: (i) a plug-and-play generator module responsible for super-resolving low-resolution Hi-C patches, (ii) a transformer-based discriminator derived from the pretrained HiCFoundation model (X. Wang *et al*., 2024), and (iii) a standardized interface that enforces consistent dimensionality and numerical representation across heterogeneous generators.

Given a low-resolution Hi-C contact matrix (40*×* 40 patches), the generator produces a super-resolved prediction that is normalized into a biologically valid interaction-frequency (IF) range and evaluated by the discriminator. The discriminator processes both generated and real high-resolution matrices to produce adversarial realism scores (Goodfellow *et al*., 2020) while also extracting intermediate feature representations used for feature-matching regularization.

The discriminator backbone uses a Vision Transformer (ViT) architecture (Dosovitskiy *et al*., 2020) pretrained on large-scale Hi-C contact maps. This design enables the adversarial classifier to encode long-range chromatin interactions, multi-scale interaction patterns, and higher-order structural dependencies that are difficult to capture using conventional convolutional discriminators.

The overall architecture of HiCP2GAN is illustrated in Fig. 1.

**Figure 1:**
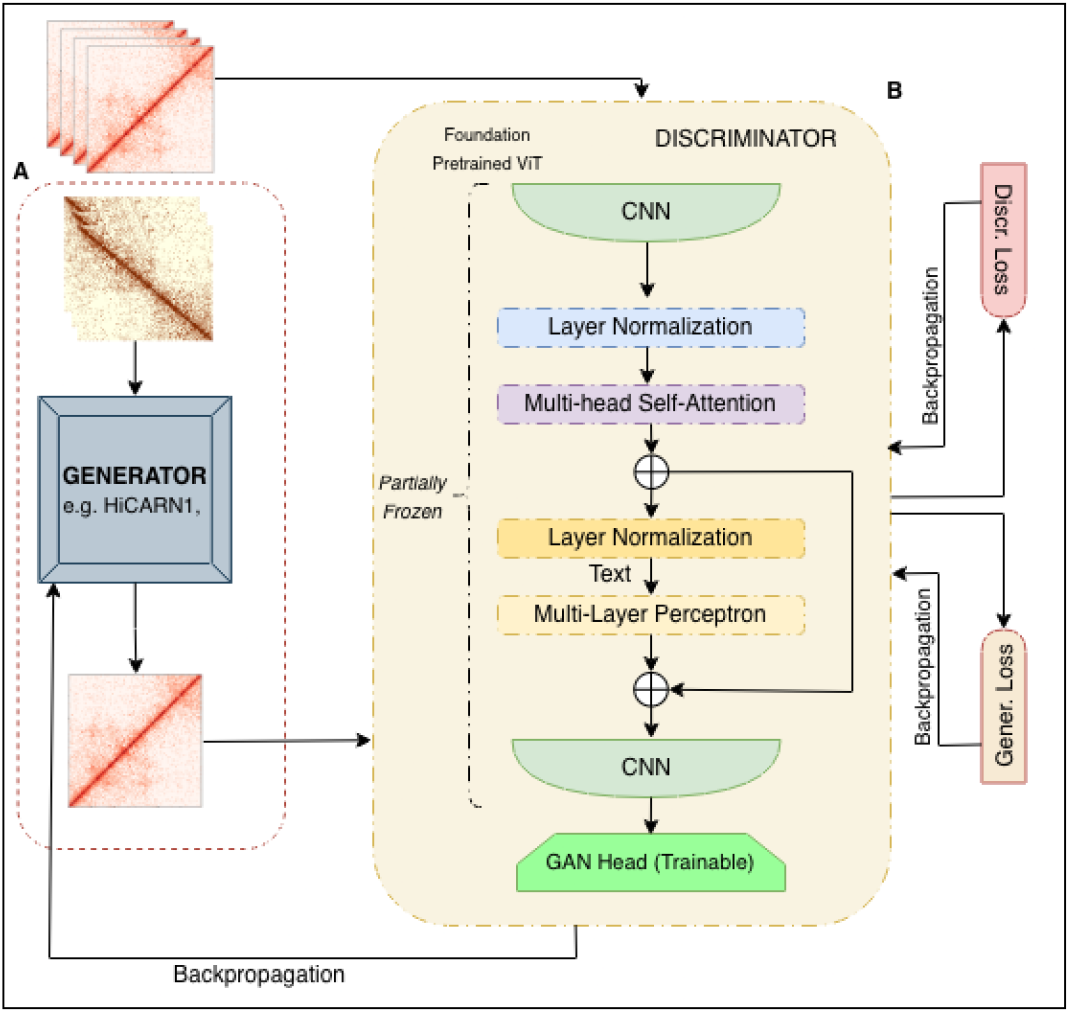
Architecture of the HiCP2GAN Plug-and-Play Adversarial Framework for Hi-C Resolution Enhancement. (A) <u>Generator</u>: A generator-agnostic super-resolution module (e.g., HiCARN1, DiCARN, or other compatible architectures) takes low-resolution Hi-C contact matrices as input and produces enhanced Hi-C maps, enabling seamless integration of diverse super-resolution models within a unified adversarial framework. (B) <u>HiCFoundation Discriminator</u>: A pretrained transformer-based discriminator initialized from HiCFoundation provides structural priors for chromatin interaction modeling. The discriminator consists of convolutional feature extractors, multi-head self-attention blocks, and feed-forward layers followed by a lightweight GAN head. The deeper transformer layers are frozen to preserve pretrained representations, while the earlier layers (closer to the input) and GAN head are finetuned during adversarial training. Adversarial and feature-matching losses guide generator optimization, enabling resolution enhancement informed by both local convolutional patterns and global chromatin interaction structure.

**Figure 2:**
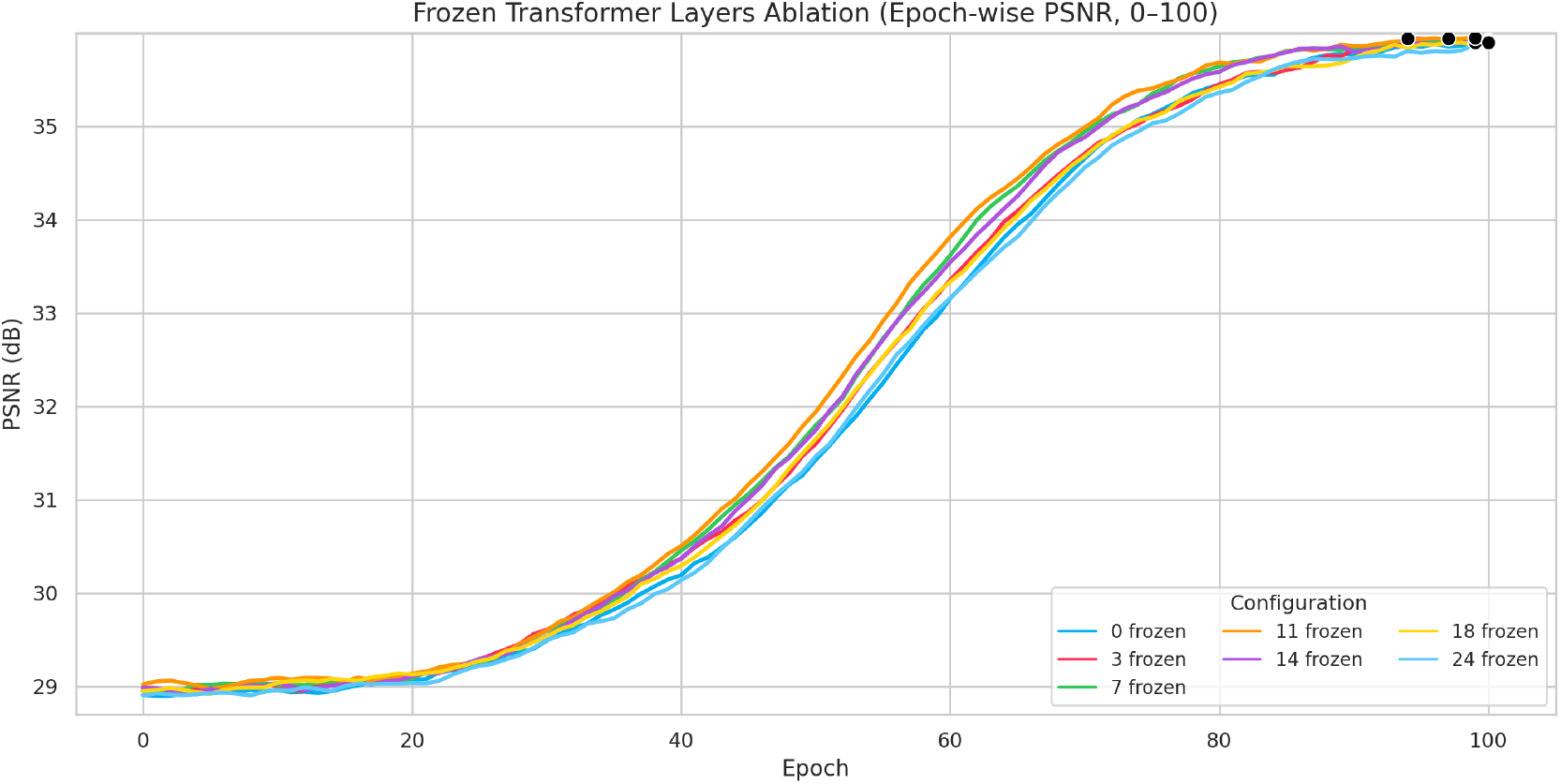
Layer finetuning validation of the HiCFoundation discriminator (100 epochs). Epoch-wise validation PSNR plots for HiCP2GAN trained with varying numbers of finetuned transformer layers (from the input side); deeper layers remain frozen. This was used to select the discriminator configuration adopted in our subsequent experiments.

### 2.2 Generator-Agnostic Interface and Data Standardization

Hi-C super-resolution models differ substantially in their architectural assumptions, including input resolutions, output ranges, preprocessing pipelines, and training objectives. These inconsistencies complicate direct comparison across methods and can introduce confounding factors during adversarial training.

HiCP2GAN addresses this problem through a generator-agnostic plug-and-play interface that standardizes input resolution and numerical representation while preserving the internal structure of each generator.

All generators operate on 40*×* 40 LR Hi-C patches and produce enhanced outputs at the same resolution. Models originally designed for 28*×* 28 inputs are integrated through deterministic resolution adaptation. Specifically, low-resolution patches are resized from 40 *×* 40 to 28*×* 28 using area-based interpolation prior to inference and restored to the original resolution using nearest-neighbor interpolation after super-resolution. No additional learnable upsampling layers are introduced, ensuring that generators are not granted extra trainable capacity beyond their original design.

To maintain numerical consistency across models, all generator outputs are normalized to the interval [0, 1]. Outputs produced using tanh activation (e.g., HiCSR) are linearly rescaled to this range, whereas outputs already operating within the target interval remain unchanged. All predictions are subsequently clamped to ensure numerical stability. Ground-truth high-resolution matrices are likewise clamped prior to computing reconstruction losses and evaluation metrics.

This standardization guarantees compatibility with reconstruction losses, perceptual similarity metrics, and biological reproducibility measures such as GenomeDISCO (Ursu *et al*., 2018), enabling unbiased benchmarking across heterogeneous generator architectures.

### 2.3 Foundation-Based Discriminator Architecture

The HiCP2GAN discriminator is derived from HiCFoundation (X. Wang *et al*., 2024), a Vision Transformer pretrained on large-scale Hi-C datasets. Input Hi-C patches are first projected to three channels through a lightweight convolutional adapter before being processed by the transformer encoder.

To adapt the pretrained encoder for adversarial learning, a lightweight fully connected classification head is appended to produce a scalar realism score indicating whether an input matrix is real or generated. During adversarial training, the deeper transformer blocks are frozen to preserve pretrained chromatin interaction representations, while the earlier blocks (towards the input end) and the adversarial head are both finetuned. The optimal finetuning depth is determined experimentally (Section 3.2).

The discriminator serves a dual role during training. In addition to adversarial classification, intermediate transformer representations are exposed for feature-matching regularization. These features enable the generator to align its outputs with structural representations learned by the pretrained foundation model, encouraging consistency with biologically meaningful chromatin interaction patterns.

### 2.4 Adversarial Objective and Optimization

HiCP2GAN is trained using a composite adversarial objective that balances structural realism, signal fidelity, and representation-level consistency across embeddings. Let *X*_*LR*_ denote a low-resolution Hi-C patch, *X*_*HR*_ the corresponding high-resolution matrix, and 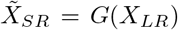 the normalized generator output.

The discriminator *D*(*·*) is trained to distinguish real high-resolution matrices from generated samples using binary cross-entropy with logits:

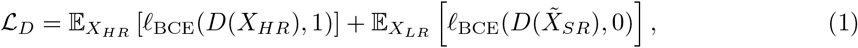

where *𝓁*_BCE_(*·,·*) denotes the binary cross-entropy loss with logits.

The generator adversarial loss encourages generated samples to be classified as real:

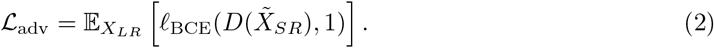

To preserve contact-intensity fidelity, we employ an *𝓁*_1_ reconstruction loss:

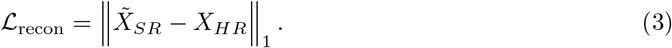

To align generated and real samples in the discriminator representation space, we further apply a feature-matching loss. Let *ϕ*(*·*) denote the pooled transformer representation extracted from the discriminator. The feature-matching loss is defined as:

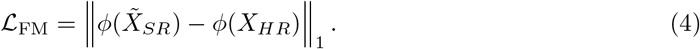

The total generator objective is:

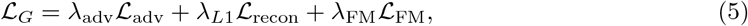

where *λ*_adv_, *λ*_*L*1_, and *λ*_FM_ are scalar weighting coefficients fixed across all experiments.

Training proceeds via alternating updates of the discriminator and generator. For each mini-batch, the discriminator is first updated using real and generated samples, followed by a generator update using the composite objective above. Both networks are optimized using the Adam optimizer with identical learning rates and momentum parameters.

Automatic mixed-precision (AMP) training is employed to improve computational efficiency while maintaining numerical stability. Gradient clipping is applied during both generator and discriminator updates to mitigate gradient explosions during early training stages.

## 3 Results

### 3.1 Experimental Setup

#### 3.1.1 Hi-C Data Description and Preprocessing

All experiments were conducted on genome-wide Hi-C contact matrices generated at a fixed resolution of 10 kb across all evaluated cell lines. The dataset includes four widely used human cell lines originating from an earlier research conducted by (Rao *et al*., 2014): GM12878 (lymphoblastoid cells), K562 (leukemia cells), HMEC (microvascular endothelial cells), and NHEK (epidermal keratinocytes).

To simulate shallow-sequenced Hi-C data, we uniformly downsampled the *in-situ* high-resolution contact maps using a ratio of 16 and 64 for a dual LR-variability study. This downsampling protocol was applied consistently across training, fine-tuning, validation, and testing datasets for all corresponding procedures.

To prevent information leakage across genomic loci, we adopt a strict chromosome-disjoint split strategy across train, validation, and test datasets as follows:

- Training chromosomes: chr1, chr3, chr5, chr7, chr8, chr9, chr11, chr13, chr15, chr17, chr18, chr19, chr21, chr22
- Validation chromosomes: chr2, chr6, chr10, chr12
- Test chromosomes: chr4, chr14, chr16, chr20

#### 3.1.2 Training

All models were trained for a fixed number of epochs with a batch size of 64, and automatic mixed precision (AMP) (Dörrich *et al*., 2023) was used to improve computational efficiency.

For our adversarial framework, the generator was optimized using a weighted combination of three loss components: (i) adversarial loss (Chen *et al*., 2020), (ii) pixel-wise *𝓁*_1_ reconstruction loss (Isola *et al*., 2017), and (iii) feature-matching loss computed from intermediate discriminator representations (Salimans *et al*., 2016). Gradient clipping was applied to both generator and discriminator updates to stabilize training.

All experiments were executed on NVIDIA TITAN RTX GPUs, while the core training and parameter optimization settings for HiCP2GAN experiments are provided in Supplementary Table S1 and Figure S1.

#### 3.1.3 Validation Protocol and Metrics

We adopted the Peak Signal-to-Noise Ratio (PSNR) (Hore and Ziou, 2010) computation metric, popularly used for measuring reconstruction fidelity for our validation benchmarking, while the Structural Similarity Index (SSIM) (Ndajah *et al*., 2010), capturing perceptual and structural consistency, and GenomeDISCO (Ursu *et al*., 2018) for assessing biological reproducibility of chromatin interaction patterns, are reported in the test scenarios.

### 3.2 Discriminator Freezing and Model Validation

We conducted a discriminator finetuning-depth ablation to characterize how the pretrained HiC-Foundation backbone responds to varying degrees of task-specific adaptation. Rather than identifying a single optimal configuration, this analysis was designed to assess the transferability of pretrained chromatin representations under adversarial supervision.

#### 3.2.1 Phase I: Discriminator Finetuning Depth Ablation

Partial layer freezing is a transfer-learning technique that preserves general features learned during large-scale pretraining while enabling task-specific adaptation (He *et al*., 2016). Howard and Ruder (2018) further demonstrated that gradual unfreezing can improve stability while preventing catastrophic forgetting. We adopt a complementary strategy: finetuning early transformer layers while freezing deeper ones, preserving higher-level pretrained representations while allowing input-proximal layers to adapt to the enhancement task. We varied the number of finetuned layers incrementally from 0 (fully frozen) to 24 (fully finetuned).

As shown in Supplementary Table S2, remarkable performance is recorded across the full range of finetuning depths, with the configurations achieving near-identical validation PSNR. Notably, the fully frozen discriminator (0 finetuned layers) performs comparably to fully or partially fine-tuned variants, suggesting that the HiCFoundation backbone transfers rich chromatin interaction representations directly to adversarial Hi-C enhancement while preserving strong performance across varying adaptation settings.

This finding strengthens the central motivation of HiCP2GAN: that a foundation-pretrained discriminator provides an inherently richer supervisory signal than a convolutional discriminator trained from scratch, regardless of finetuning configuration. Given this plateau, the objective of discriminator configuration selection shifts from maximizing reconstruction fidelity to identifying the most parameter-efficient operating point within the stable performance region. The best-performing configurations clustered between 10 and 17 finetuned layers, which we carry forward into Phase II for efficiency-targeted analysis.

Extended training (500 epochs) yields modest gains (Supplementary Table S3) but the additional compute overhead is not justified; hence, we adopt 100 epochs for all experiments.

#### 3.2.2 Phase II: Depth Efficiency-Targeted Discriminator Fine-Tuning

To further refine the fidelity capability recorded in phase I, we conducted a second-phase experimentation solely focusing on this high-performing configuration cluster. In this phase, each configuration within the range of 10–17 blocks was trained independently with all selected layers fully fine-tuned while the deeper layers were completely truncated. This phase was inspired by the need to identify the most stable and efficient operating point within the optimal depth region identified during Phase I.

Table 2 reports the resulting PSNR values averaged across runs with their corresponding statistical variability. The results confirm that performance remains remarkably stable across the cluster, with minimal variance across configurations. However, the layout involving fine-tuning the first 11 transformer layers provided the best trade-off between reconstruction quality and training efficiency, achieving the highest average PSNR and best standard deviation. Hence, the adoption as the prefered HiCP2GAN discriminator configuration.

**Table II:** Phase II efficiency-targeted finetuning within the 10–17 layer cluster on GM12878. Each configuration is trained independently with the selected layers fully finetuned and deeper layers discarded. HiCARN1 is used as the generator. PSNR is reported as mean*±* standard deviation across runs.

| Fine-tuned Layers | PSNR (dB) ± SD |
| --- | --- |
| 10 | 35.93 ± 0.003 |
| 11 | **35.93 ± 0.002** |
| 12 | 35.92 ± 0.003 |
| 13 | 35.93 ± 0.003 |
| 14 | 35.92 ± 0.004 |
| 15 | 35.92 ± 0.003 |
| 16 | 35.93 ± 0.004 |
| 17 | 35.92 ± 0.003 |

### 3.3 HiCP2GAN Improves Hi-C Resolution Over Standalone Generators

Table III compares standalone generators with their HiCP2GAN adaptation. Every evaluated generator, including DiCARN, HiCARN1, HiCNN, and HiCPlus, improves when trained within the HiCP2GAN framework. The gains are largely consistent across PSNR, SSIM, and GenomeDISCO, reinforcing the hypothesis that foundation-supervised adversarial learning benefits a range of architectures training and testing on the same cell line (GM12878), but mutually exclusive chromosomes.

**Table III:** Same-cell evaluation on GM12878. Standalone generators, the respective HiCP2GAN-integrated variants, and traditional GAN-based models are compared on chromosomes 4, 14, 16, and 20 of the GM12878 cell line. Bold values indicate improvement over standalone, underline indicates the best traditional GAN-based method, and (*) denotes the best-performing method overall for each metric.

| Generator | Standalone |  |  | + HiCP2GAN |  |  | Method | Traditional GAN-based |  |  |
| --- | --- | --- | --- | --- | --- | --- | --- | --- | --- | --- |
|  | PSNR | SSIM | GenomeDISCO | PSNR | SSIM | GenomeDISCO |  | PSNR | SSIM | GenomeDISCO |
| DiCARN | 35.72 | 0.9142 | 0.9138 | **36.02\*** | **0.9213\*** | **0.9196\*** | EnHiC | 28.9410 | 0.7917 | 0.8484 |
| HiCARN1 | 35.64 | 0.9138 | 0.9106 | **35.93** | **0.9202** | **0.9140** | hicGAN | 32.0452 | 0.8501 | 0.8613 |
| HiCNN | 35.46 | 0.9085 | 0.9079 | **35.84** | **0.9161** | **0.9147** | HiCARN2 | <u>35.2353</u> | <u>0.9128</u> | <u>0.9174</u> |
| HiCPlus | 35.32 | 0.9051 | 0.9044 | **35.70** | **0.9124** | **0.9103** | DeepHiC | 34.8090 | 0.9053 | 0.9105 |

### 3.4 Generalizing HiCP2GAN Across Unseen Cell Lines

To assess whether the fidelity restoration capability of HiCP2GAN transfers to cell lines not seen during training, we evaluated models trained only on GM12878 on three other human cell lines: HMEC, K562, and NHEK. Tables IV, V, and VI report per-cell-line comparisons of each generator in its standalone form against the same generator integrated with HiCP2GAN.

**Table IV:** Cross-cell evaluation on the K562 cell line. Models trained on GM12878 are evaluated on chromosomes 4, 14, 16, and 20. Bold values indicate the better result between standalone and HiCP2GAN-augmented versions of the same generator. The best traditional GAN-based method is underlined, while (*) denotes the best-performing method overall for each metric.

| Generator | Standalone |  |  | + HiCP2GAN |  |  | Traditional GAN-based |  |  |  |
| --- | --- | --- | --- | --- | --- | --- | --- | --- | --- | --- |
|  | PSNR | SSIM | GenomeDISCO | PSNR | SSIM | GenomeDISCO | Method | PSNR | SSIM | GenomeDISCO |
| DiCARN | 33.89 | 0.9450 | 0.8319 | **34.53** | **0.9477\*** | **0.8449\*** | EnHiC | 24.7901 | 0.7193 | 0.7558 |
| HiCARN1 | 33.77 | 0.9319 | 0.8110 | **34.68\*** | **0.9446** | **0.8316** | hicGAN | 31.8010 | 0.8107 | 0.7670 |
| HiCNN | 31.40 | 0.8535 | **0.7958** | **31.73** | **0.8588** | 0.7060 | HiCARN2 | 34.3449 | 0.9472 | 0.8327 |
| HiCPlus | 31.57 | 0.8236 | 0.7627 | **31.81** | **0.8623** | **0.7702** | HiCSR | 16.1466 | 0.3168 | 0.7639 |

**Table V:** Cross-cell evaluation on the HMEC cell line. Models trained on GM12878 are evaluated on chromosomes 4, 14, 16, and 20. Bold values indicate the better result between standalone and HiCP2GAN-augmented versions of the same generator. The best traditional GAN-based method is underlined, while (*) denotes the best-performing method overall for each metric.

| Generator | Standalone |  |  | + HiCP2GAN |  |  | Traditional GAN-based |  |  |  |
| --- | --- | --- | --- | --- | --- | --- | --- | --- | --- | --- |
|  | PSNR | SSIM | GenomeDISCO | PSNR | SSIM | GenomeDISCO | Method | PSNR | SSIM | GenomeDISCO |
| DiCARN | 32.58 | 0.8628 | 0.5171 | **34.12** | **0.8869** | **0.5214** | EnHiC | 26.76 | 0.6416 | 0.4425 |
| HiCARN1 | 33.99 | **0.8816** | 0.5027 | **34.09** | 0.8711 | **0.5819\*** | hicGAN | 31.13 | 0.8807 | 0.4371 |
| HiCNN | 26.66 | 0.8602 | 0.5011 | **27.92** | **0.8794** | **0.5030** | HiCARN2 | <u>35.06*</u> | <u>0.9741*</u> | 0.5185 |
| HiCPlus | 26.11 | 0.7577 | 0.4445 | **29.41** | **0.7897** | **0.4611** | HiCSR | 16.07 | 0.3971 | 0.4448 |

**Table VI:** Cross-cell evaluation on the NHEK cell line. Models trained on GM12878 are evaluated on chromosomes 4, 14, 16, and 20. Bold values indicate the better result between standalone and HiCP2GAN-augmented versions of the same generator. The best traditional GAN-based method is underlined, while (*) denotes the best-performing method overall for each metric.

| Generator | Standalone |  |  | + HiCP2GAN |  |  | Traditional GAN-based |  |  |  |
| --- | --- | --- | --- | --- | --- | --- | --- | --- | --- | --- |
|  | PSNR | SSIM | GenomeDISCO | PSNR | SSIM | GenomeDISCO | Method | PSNR | SSIM | GenomeDISCO |
| DiCARN | 35.33 | 0.9636 | 0.8493 | **35.58\*** | **0.9655** | **0.8547\*** | EnHiC | 25.39 | 0.6589 | 0.5662 |
| HiCARN1 | 35.28 | 0.9541 | 0.8163 | **35.55** | **0.9681** | **0.8208** | hicGAN | 31.39 | 0.8250 | 0.5743 |
| HiCNN | 34.13 | 0.8830 | 0.8178 | **35.13** | **0.9091** | **0.8178** | HiCARN2 | <u>35.08</u> | <u>0.9712*</u> | 0.6497 |
| HiCPlus | 33.15 | 0.8491 | **0.8384** | **34.02** | **0.8689** | 0.7987 | HiCSR | 16.08 | 0.255 | 0.5689 |

For each generator, we report performance in two settings: (i) standalone and (ii) augmented with HiCP2GAN (Standalone + HiCP2GAN). Bold values indicate the better result between the standalone and augmented versions of the same generator, highlighting the effect of the proposed plug-and-play framework. The best traditional GAN is underlined, while (*) denotes the overall best method for each metric, enabling direct comparison between augmented models and traditional GAN-based approaches. Across unseen cell lines, the foundation-based discriminator enables generators to maintain or improve structural and intensity fidelity. For generators that already perform well standalone (e.g., DiCARN and HiCARN1), HiCP2GAN preserves high PSNR and SSIM under a shift in distribution; where standalone performance is limited, pairing with HiCP2GAN often yields gains in at least one of PSNR, SSIM, or GenomeDISCO. HiCARN1 paired with HiCP2GAN achieves strong cross-cell results (Tables IV–VI), consistent with its same-cell behaviour.

Overall, the plug-and-play design generalizes: a single training setup on GM12878 transfers to multiple unseen cell lines without cell-specific retraining. Per-chromosome PSNR, SSIM, and GenomeDISCO on the same test chromosomes are reported in Supplementary Table S4.

### 3.5 HiCP2GAN Improves Performance Across Seen and Unseen Cell Lines

The comparative results in Tables III–VI show that HiCP2GAN consistently improves generator performance across both same-cell (GM12878) and cross-cell (HMEC, K562, NHEK) settings.

On the same-cell setting (Table III), HiCP2GAN achieves the best overall performance across all evaluated metrics, outperforming both standalone generators and traditional GAN-based methods. In particular, the HiCP2GAN-integrated DiCARN model achieves the highest PSNR, SSIM, and GenomeDISCO, indicating improved reconstruction quality and structural fidelity. When compared to traditional GAN-based methods, including strong baselines such as HiCARN2, HiCP2GAN remains competitive across computational metrics and often achieves improved biological consistency, as reflected by GenomeDISCO. Underline indicates the best-performing traditional GAN-based method, while (*) denotes the overall best method for each metric. Notably, HiCP2GAN-augmented models frequently achieve the overall best results, although performance varies across cell lines and metrics.

Importantly, these trends persist under cross-cell evaluation (Tables III, IV, V, and VI), demonstrating that the gains introduced by HiCP2GAN generalize beyond same-cell settings. While not uniformly dominant across all unseen cell lines, HiCP2GAN consistently enhances generator performance and maintains strong structural fidelity under distribution shift.

### 3.6 HiCP2GAN Improves Loop Detection Over Standalone Generators

Fine-scale chromatin loops are fundamental structural units of 3D genome organization, linking regulatory elements through long-range interactions. Accurate loop detection from enhanced Hi-C data therefore provides an important downstream validation of enhancement quality beyond pixel-level reconstruction metrics.

To assess biological loop recovery, we called loops from enhanced contact maps on GM12878 test chromosomes (chr4, chr14, chr16, and chr20) using Fit-Hi-C (Kaul *et al*., 2020) and compared them with loops detected from the corresponding ground-truth 10kb HR matrices. Recovery was quantified using the F1 score, which balances precision and recall.

Figure 3 summarizes the results. Across all evaluated generators (HiCNN, DiCARN, HiCARN1, and HiCPlus), HiCP2GAN-integrated models outperform their standalone variants, indicating that the improvements in contact-map quality translates into improved downstream structural inference. This is exemplified in HiCP2GAN-DiCARN’s performance, having a higher overall F1 score of 0.8295 compared with 0.8078 for standalone DiCARN, and this pattern is observed to be consistent across the other evaluated generators, showing that the benefit of HiCP2GAN is not architecture-specific.

**Figure 3:**
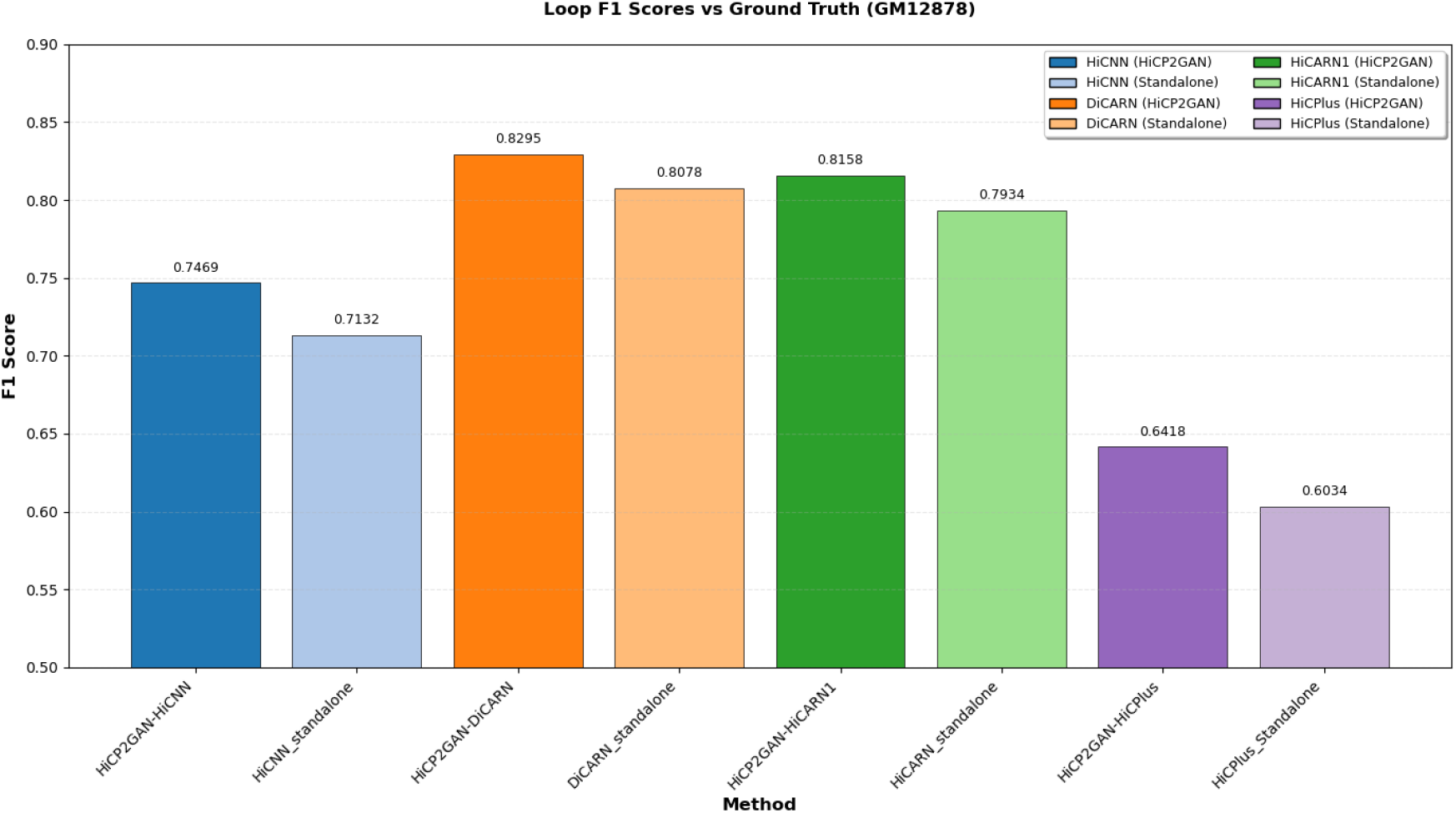
Loop F1 scores from test chromosomes (chr4, chr14, chr16, chr20) compared to ground truth on GM12878. Dark bars represent HiCP2GAN-integrated models and light bars represent standalone generators. HiCP2GAN consistently improves loop recovery, with HiCP2GAN-DiCARN achieving the highest F1 score.

### 3.7 HiCP2GAN Improves TAD Detection and Quality

Topologically associating domains (TADs) are fundamental organizational units of chromatin architecture, delineating self-interraction genomic regions where contacts are significantly enriched (Dixon *et al*., 2012). Functions ranging enhancer-promoted facilitation to gene regulation are critically supported by this compartmentalization concept of functional genomics, especially since the disruption of TAD boundaries has been linked to annomalies in gene activation and diseases (Dixon *et al*., 2012). A model’s capacity to recover TADs from enhanced Hi-C data therefore serves as a biologically meaningful indicator of its utility beyond numerical reconstruction metrics.

To assess TAD recovery, we applied TopDom (Shin *et al*., 2016) to detect TADs from the 60 kb to 2.45 Mb region of chromosome 14 in the K562 cell line, using both enhanced contact maps produced by each method and the ground-truth high-resolution Hi-C data. We then use the Jaccard similarity coefficent (Besta *et al*., 2020), to quantify similarity concordance in TAD recovered across the models. Higher Jaccard scores indicate better boundary recovery.

Table VII reports Jaccard scores for each generator in both standalone and HiCP2GAN-based configurations, while Fig. 4 provides qualitative boundary comparisons. HiCP2GAN-integrated models are observed to consistently recover TAD boundaries with higher concordance to the ground truth across all evaluated architectures.

**Table VII:** TAD boundary recovery on K562 chromosome 14. Jaccard similarity (%) between TAD boundaries detected by TopDom on imputed contact maps and those identified in the ground-truth HR Hi-C data, evaluated over the 60 kb–2.45 Mb region. Higher scores indicate more consistent boundary recovery. Bold represents higher recovery.

| Generator | Standalone (%) | + HiCP2GAN (%) |
| --- | --- | --- |
| DiCARN | 55.91 | **61.28** |
| HiCARN1 | 56.15 | **61.01** |
| HiCNN | 41.51 | **43.88** |
| HiCPlus | 40.71 | **41.82** |

**Figure 4:**
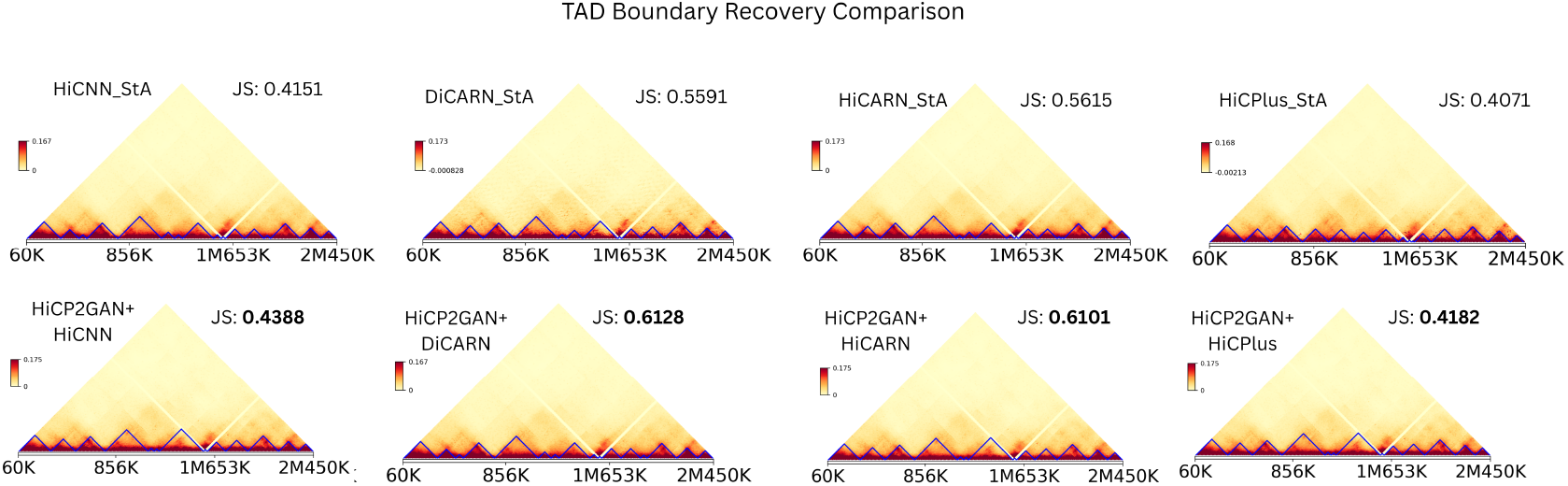
TAD structure comparison on K562 chromosome 14 (60 kb – 2.45 Mb). Contact maps and TopDom-detected TAD boundaries for each generator in standalone and HiCP2GAN-integrated configurations, alongside the ground-truth high-resolution Hi-C map. HiCP2GAN-enhanced maps exhibit boundary patterns more consistent with the ground truth, as quantified by the Jaccard scores (JS) labelled in the images, as well as represented in Table VII.

### 3.8 HiCP2GAN Improves 3D Chromosome Reconstruction

Beyond contact-map fidelity, we evaluated whether HiCP2GAN-enhanced matrices yield more accurate 3D chromosome structures. We reconstructed chromatin conformations from enhanced Hi-C contact maps using 3DUnicorn (Chandrashekar *et al*., 2025) and compared them to the reference structures derived from high-resolution data using the Spearman correlation coefficient (SCC) (Ali Abd Al-Hameed, 2022) over the 18.5Mbp–22.5Mbp region of chromosome 4 in the GM12878 cell line. Figure 5 shows 3D reconstructions for four generators in both standalone and HiCP2GAN-integrated settings, where the light green chain represents the reference structure and the colored chain denotes the predicted structure. HiCP2GAN improves SCC for all four generators, with the largest gains observed for the weaker standalone baselines (HiCPlus and HiCNN), and HiCP2GAN + DiCARN attaining the highest SCC (0.9896). Overall, HiCP2GAN consistently improves 3D reconstruction quality, establishing enhanced biological relevance by our proposed method.

**Figure 5:**
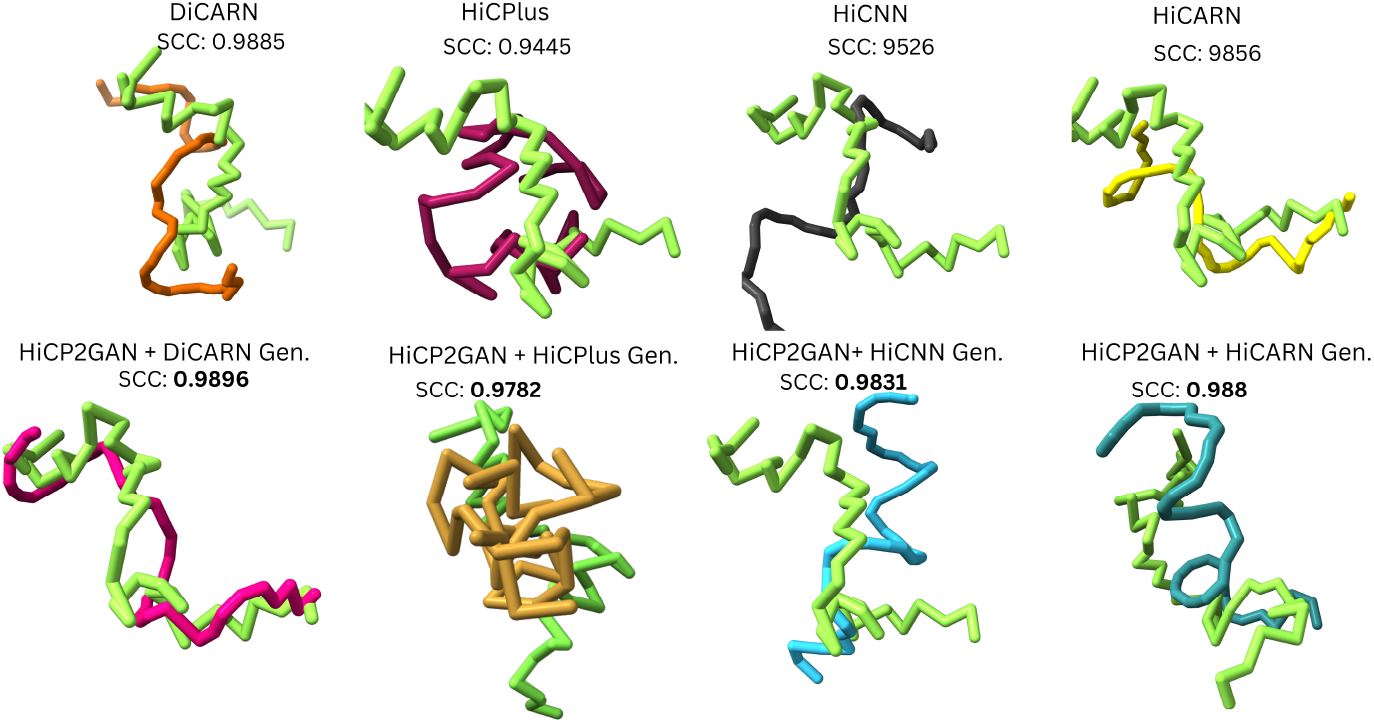
Standalone Generators vs HiCP2GAN Variants: 3D chromosome reconstruction. Each row compares a generator alone (top) with HiCP2GAN + generator (bottom). Light green: reference structure; colored chain: predicted structure. SCC: Spearman correlation coefficient.

## 4 Discussion and Conclusion

HiCP2GAN introduces a plug-and-play adversarial framework for Hi-C resolution enhancement that leverages a pretrained Hi-C foundation model as its discriminator. By decoupling generator design from adversarial supervision, the framework enables consistent evaluation across diverse super-resolution architectures while benefiting from foundation-level representations. Ablation studies show that the pretrained HiCFoundation backbone transfers effectively to adversarial Hi-C enhancement with minimal task-specific adaptation, maintaining stable performance across finetun-ing configurations. Pretraining on large-scale Hi-C data provides an inductive bias that allows the discriminator to guide generators toward biologically meaningful reconstructions. The efficiency-targeted Phase II analysis further identifies a compact configuration that preserves this capability without requiring full model complexity. Importantly, the gains produced by HiCP2GAN extend beyond pixel-level reconstruction. Improvements in loop detection F1 and TAD boundary recovery indicate that foundation-supervised adversarial training enhances downstream structural inference. The observed inverse relationship between standalone generator strength and HiCP2GAN gains suggests that the framework is particularly beneficial for architectures that struggle to preserve local chromatin structure.

Overall, HiCP2GAN shifts the focus of Hi-C reconstruction from architecture-specific optimization to shared adversarial supervision and representational alignment. Its generator-agnostic design, coupled with robust cross-cell generalization, positions it both as a benchmarking framework and as a generalizable training scaffold for future Hi-C enhancement methods.

## 5 Author contributions statement

S.O. designed the pipeline, wrote the code, and wrote the initial draft manuscript. S.O. and O.O. analyzed the results. O.O. conceived and supervised the project. All authors wrote and reviewed the manuscript.

## 6 Acknowledgements

## 7 Code and Data Availability

HiCP2GAN is a containerized software made available via: https://github.com/OluwadareLab/HiCP2GAN. The Hi-C datasets used in this study including GM12878, K562, HMEC, and NHEK (Rao *et al*., 2014) GEO Accession Database via GEO code GSE63525. Pretrained and trained model weights are also made available via https://zenodo.org/records/20030290.

## 8 Supplemental Data

Supplementary figures and tables are included in the Supplementary Materials document.

## 9 Competing interests

No competing interest is declared.

## 10 Funding

This work was supported by the National Institutes of General Medical Sciences of the National Institutes of Health under award number R35GM150402 to O.O.

## Notes

### Competing Interest Statement

The authors have declared no competing interest.

## References

Ali Abd Al-Hameed, Khawla (2022). “Spearman’s correlation coefficient in statistical analysis”, International Journal of Nonlinear Analysis and Applications, Vol. 13 No. 1, pp. 3249–3255.

Besta, Maciej et al., (2020). “Communication-efficient jaccard similarity for high-performance distributed genome comparisons”, 2020 IEEE International Parallel and Distributed Processing Symposium (IPDPS). IEEE, pp. 1122–1132.

Chandrashekar, Mohan Kumar B et al., (2025). “Unicorn: enhancing single-cell Hi-C data with blind super-resolution for 3D genome structure reconstruction”, Bioinformatics, Vol. 41 No. Supplement 1, pp. i475–i483.

Chen, Minghao et al., (2020). “Adversarial-learned loss for domain adaptation”, Proceedings of the AAAI conference on artificial intelligence. Vol. 34. No. 04, pp. 3521–3528.

Dimmick, Michael (2020). HiCSR: a Hi-C super-resolution framework for producing highly realistic c University of Toronto (Canada).

Dixon, Jesse R et al., (2012). “Topological domains in mammalian genomes identified by analysis of chromatin interactions”, Nature, Vol. 485 No. 7398, pp. 376–380.

Dörrich, Marion, Fan, Mingcheng, and Kist, Andreas M (2023). “Impact of mixed precision techniques on training and inference efficiency of deep neural networks”, IEEE Access, Vol. 11, pp. 57627–57634.

Dosovitskiy, Alexey et al., (2020). “An image is worth 16 × 16 words: Transformers for image recognition at scale”, arXiv preprint arXiv:2010.11929,

Goodfellow, Ian et al., (2020). “Generative adversarial networks”, Communications of the ACM, Vol. 63 No. 11, pp. 139–144.

He, Kaiming et al., (2016). “Deep residual learning for image recognition”, Proceedings of the IEEE conference on computer vision and pattern recognition, pp. 770–778.

Hicks, Parker and Oluwadare, Oluwatosin (2022). “HiCARN: resolution enhancement of Hi-C data using cascading residual networks”, Bioinformatics, Vol. 38 No. 9, pp. 2414–2421.

Hong, Hao et al., (2020). “DeepHiC: A generative adversarial network for enhancing Hi-C data resolution”, PLoS computational biology, Vol. 16 No. 2, e1007287.

Hore, Alain and Ziou, Djemel (2010). “Image quality metrics: PSNR vs. SSIM”, 2010 20th international conference on pattern recognition. IEEE, pp. 2366–2369.

Howard, Jeremy and Ruder, Sebastian (2018). “Universal language model fine-tuning for text classification”, Proceedings of the 56th Annual Meeting of the Association for Computational Linguist pp. 328–339.

Hu, Yangyang and Ma, Wenxiu (2021). “EnHiC: learning fine-resolution Hi-C contact maps using a generative adversarial framework”, Bioinformatics, Vol. 37 No. Supplement 1, pp. i272–i279.

Imakaev, Maxim et al., (2012). “Iterative correction of Hi-C data reveals hallmarks of chromosome organization”, Nature methods, Vol. 9 No. 10, pp. 999–1003.

Isola, Phillip et al., (2017). “Image-to-image translation with conditional adversarial networks”, Proceedings of the IEEE conference on computer vision and pattern recognition, pp. 1125–1134.

Kaul, Arya, Bhattacharyya, Sourya, and Ay, Ferhat (2020). “Identifying statistically significant chromatin contacts from Hi-C data with FitHiC2”, Nature protocols, Vol. 15 No. 3, pp. 991–1012.

Li, Hao et al., (2025). “Foundir: Unleashing million-scale training data to advance foundation models for image restoration”, Proceedings of the IEEE/CVF international conference on computer vision, pp. 12626–12636.

Li, Pingjing et al., (2025). “HiADN: Lightweight Resolution Enhancement of Hi-C Data Using High Information Attention Distillation Network”, IEEE Transactions on Computational Biology and Bioinformatics,

Li, Zhilan and Dai, Zhiming (2020). “SRHiC: a deep learning model to enhance the resolution of Hi-C data”, Frontiers in genetics, Vol. 11, p. 519766.

Lieberman-Aiden, Erez et al., (2009). “Comprehensive mapping of long-range interactions reveals folding principles of the human genome”, science, Vol. 326 No. 5950, pp. 289–293.

Liu, Qiao, Lv, Hairong, and Jiang, Rui (2019). “hicGAN infers super resolution Hi-C data with generative adversarial networks”, Bioinformatics, Vol. 35 No. 14, pp. i99–i107.

Liu, Tong and Wang, Zheng (2019a). “HiCNN: a very deep convolutional neural network to better enhance the resolution of Hi-C data”, Bioinformatics, Vol. 35 No. 21, pp. 4222–4228.

Liu, Tong and Wang, Zheng (2019b). “HiCNN2: enhancing the resolution of Hi-C data using an ensemble of convolutional neural networks”, Genes, Vol. 10 No. 11, p. 862.

Ma, Chenxi et al., (2024). “Pretraining a foundation model for generalizable fluorescence microscopy-based image restoration”, Nature Methods, Vol. 21 No. 8, pp. 1558–1567.

Ndajah, Peter et al., (2010). “SSIM image quality metric for denoised images”, Proc. 3rd WSEAS Int. Conf. on Visualization, Imaging and Simulation. Vol. 53.

Olowofila, Samuel and Oluwadare, Oluwatosin (2025). “DiCARN-DNase: enhancing cell-to-cell Hi-C resolution using dilated cascading ResNet with self-attention and DNase-seq chromatin accessibility data”, Bioinformatics, Vol. 41 No. 9, btaf452.

Rao, Suhas SP et al., (2014). “A 3D map of the human genome at kilobase resolution reveals principles of chromatin looping”, Cell, Vol. 159 No. 7, pp. 1665–1680.

Salimans, Tim et al., (2016). “Improved techniques for training gans”, Advances in neural information processing systems, Vol. 29.

Shin, Hanjun et al., (2016). “TopDom: an efficient and deterministic method for identifying topological domains in genomes”, Nucleic acids research, Vol. 44 No. 7, e70–e70.

Tej, Akella Ravi et al., (2020). “Enhancing perceptual loss with adversarial feature matching for super-resolution”, 2020 International joint conference on neural networks (IJCNN). IEEE, pp. 1–8.

Ursu, Oana et al., (2018). “GenomeDISCO: a concordance score for chromosome conformation capture experiments using random walks on contact map graphs”, Bioinformatics, Vol. 34 No. 16, pp. 2701–2707.

Vaswani, Ashish et al., (2017). “Attention is all you need”, Advances in neural information processing systems, Vol. 30.

Wang, Xiao et al., (2024). “A generalizable Hi-C foundation model for chromatin architecture, single-cell and multi-omics analysis across species”, bioRxiv,

Zhang, Yan et al., (2018). “Enhancing Hi-C data resolution with deep convolutional neural network HiCPlus”, Nature communications, Vol. 9 No. 1,. 750.

