## Supplemental File 1 for "HiCP2GAN: A Plug and Play Foundation Model-based GAN for Hi-C Enhancement"

This supplementary document records implementation details, auxiliary benchmarks, and extended tables referenced from the main manuscript. Unless noted otherwise, all HiCP2GAN runs use the HiCFoundation ViT discriminator described in the main text from Wang et al. [2024], the generator-agnostic interface described in the main text, and 10 kb contact maps with the chromosome-disjoint splits given in the main Methods.

#### Methods

##### Supplementary Figure S1: architecture overview

Figure S1 summarizes the data flow and losses used in HiCP2GAN; it extends the conceptual overview in the main manuscript with implementation-level detail.

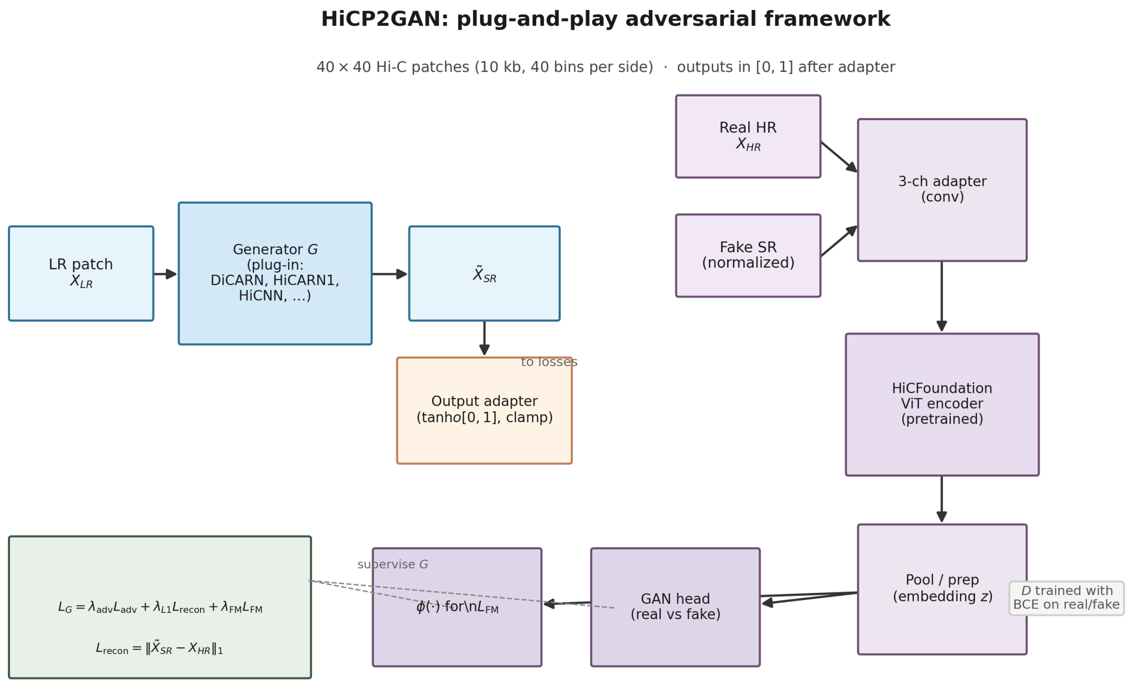


**Fig. S1. Supplementary Figure S1. Overview of HiCP2GAN.** *Left:* a plug-in generator $G$ maps a low-resolution patch $X_{LR}$ to $\tilde{X}_{SR}$, followed by the module-aware output adapter involving tanh rescaling and clamping to $[0, 1]$. *Right:* the discriminator processes real $X_{HR}$ and normalized fake $\tilde{X}_{SR}$ through a three-channel convolutional adapter and the HiCFoundation ViT encoder; pooled embeddings feed both the adversarial head (BCE) and the representation $\phi(\cdot)$ used in feature matching. *Bottom:* the composite generator objective combines adversarial, $\ell_1$ reconstruction, and feature-matching terms; the discriminator is trained alternately on real and fake inputs.

##### Supplementary Table S1: default training hyperparameters

Table S1 lists the default training and optimization settings used for the reported HiCP2GAN experiments, except where a hyperparameter study explicitly varies a quantity (e.g. discriminator depth in the freeze sweep).

##### Discriminator adaptation (HiCFoundation)

In section 3.2 of the manuscript, we describe a compact operating point with the first eleven transformer blocks finetuned and deeper blocks frozen (Phase II), following the validation study summarized there. Feature matching applies $\ell_1$ loss to the discriminator embedding used for the adversarial head (the same tensor as $\phi(\cdot)$ in the main text).

**Table S1. Supplementary Table S1.** Core training and optimization settings for HiCP2GAN experiments.

| **Setting** | **Value** |
|---|---|
| Patch size (LR / HR) | $40 \times 40$ bins |
| Batch size | 64 |
| Training epochs (main experiments) | 100 |
| Mixed precision (AMP) | Enabled |
| Optimizer | Adam, $\beta_1$=0.5, $\beta_2$=0.999, weight decay 0 |
| Learning rate — generator | $2 \times 10^{-4}$ |
| Learning rate — discriminator | $2 \times 10^{-4}$ |
| $\lambda_{\text{adv}}, \lambda_{L1}, \lambda_{\text{FM}}$ | 1, 10, 10 |
| Gradient clipping (max norm) | 1.0 (generator and discriminator) |
| Random seed | 1337 (single-seed runs unless otherwise stated) |
| ViT backbone | ViT-L/16 (HiCFoundation) |
| Discriminator MLP width | 256 |

#### Generator integration

**Resolution adaptation ($40 \to 28 \to 40$).** HiCSR, HiCNN, and HiCPlus are wrapped so that LR patches are downsampled with area interpolation to $28 \times 28$, the native generator runs at that resolution, and outputs are upsampled to $40 \times 40$ with nearest-neighbour interpolation. HiCARN1, DiCARN, and DeepHiC consume $40 \times 40$ patches directly.

**Output normalization.** HiCSR tanh outputs are mapped to $[0, 1]$ via $(x+1)/2$; all generator outputs and HR targets are clamped to $[0, 1]$ before reconstruction and GenomeDISCO computations.

#### Low-resolution downsampling protocol (1/16 vs. 1/64)

Following Rao et al. [2014] high-depth maps, low-resolution inputs are generated by uniform downsampling of contact counts to 1/16 or 1/64 of effective sequencing depth, applied consistently across train, validation, and test partitions. Primary manuscript metrics for same-cell GM12878 and cross-cell transfer use 1/16 downsampling unless a supplementary table explicitly reports 1/64.

#### Hyperparameter search

**Table S2.** Phase I discriminator finetuning-depth sweep on GM12878 (100 epochs). Average validation PSNR when the first $N$ transformer blocks of the HiCFoundation backbone are finetuned and the remaining $24-N$ deeper blocks are frozen. HiCARN1 is used as the generator throughout.

| Finetuned | PSNR | Finetuned | PSNR | Finetuned | PSNR |
|---|---|---|---|---|---|
| 0 | 35.92 | 9 | 35.92 | 18 | 35.92 |
| 1 | **35.93** | 10 | **35.93** | 19 | 35.91 |
| 2 | 35.92 | 11 | 35.92 | 20 | 35.91 |
| 3 | 35.92 | 12 | 35.91 | 21 | 35.91 |
| 4 | 35.91 | 13 | **35.93** | 22 | 35.92 |
| 5 | 35.92 | 14 | 35.92 | 23 | 35.92 |
| 6 | 35.92 | 15 | 35.92 | 24 | 35.91 |
| 7 | 35.92 | 16 | **35.93** | | |
| 8 | 35.92 | 17 | **35.93** | | |

#### Supplementary Table S3: extended discriminator training (500 epochs)

The main text cites this table when noting that 500-epoch training of the top Phase I configurations yields only marginal PSNR gains relative to the 100-epoch protocol. HiCARN1 served as the generator; metrics are mean validation PSNR on GM12878.

**Table S3.** Extended 500-epoch training for selected discriminator finetuning depths. Average validation PSNR (dB) on GM12878 with HiCARN1 as the generator.

| Finetuned transformer blocks (from input) | PSNR (dB) |
|---|---|
| 10 | 35.93 |
| 13 | **35.94** |
| 16 | 35.93 |
| 17 | 35.93 |

### Results

#### Supplementary Table S4: per-chromosome cross-cell metrics (1/16)

Table S4 lists PSNR (dB), SSIM, and GenomeDISCO on each held-out test chromosome (chr4, chr14, chr16, chr20) for models trained on GM12878 and evaluated on HMEC, K562, and NHEK (1/16 downsampling). Pooled metrics appear in the main text; this table gives the per-chromosome breakdown with cross-chromosome variation.

Table S4: Per-chromosome cross-cell evaluation (1/16 downsampling) on test chromosomes 4, 14, 16, and 20. *Note: All values are compressed to 4 decimal points.*

| | | | **Standalone** | | | **+ HiCP2GAN** | | |
|---|---|---|---|---|---|---|---|---|
| **Cell line** | **Chr** | **Generator** | PSNR | SSIM | GDISCO | PSNR | SSIM | GDISCO |
| *HMEC (cross-cell, 1/16 LR)* | | | | | | | | |
| HMEC | chr4 | DiCARN | 31.92 | 0.8571 | 0.5102 | 33.47 | 0.8823 | 0.5145 |
| HMEC | chr4 | HiCARN1 | 33.21 | 0.8765 | 0.4958 | 33.42 | 0.8665 | 0.5748 |
| HMEC | chr4 | HiCNN | 26.11 | 0.8545 | 0.4942 | 27.28 | 0.8741 | 0.4961 |
| HMEC | chr4 | HiCPlus | 25.67 | 0.7523 | 0.4380 | 28.88 | 0.7845 | 0.4548 |
| HMEC | chr14 | DiCARN | 32.44 | 0.8612 | 0.5156 | 33.89 | 0.8851 | 0.5199 |
| HMEC | chr14 | HiCARN1 | 33.78 | 0.8798 | 0.5012 | 33.91 | 0.8698 | 0.5806 |
| HMEC | chr14 | HiCNN | 26.45 | 0.8589 | 0.4998 | 27.67 | 0.8782 | 0.5018 |
| HMEC | chr14 | HiCPlus | 25.98 | 0.7567 | 0.4432 | 29.15 | 0.7889 | 0.4599 |
| HMEC | chr16 | DiCARN | 33.18 | 0.8645 | 0.5211 | 34.56 | 0.8889 | 0.5254 |
| HMEC | chr16 | HiCARN1 | 34.44 | 0.8834 | 0.5068 | 34.38 | 0.8723 | 0.5863 |
| HMEC | chr16 | HiCNN | 27.02 | 0.8618 | 0.5053 | 28.34 | 0.8809 | 0.5072 |
| HMEC | chr16 | HiCPlus | 26.34 | 0.7589 | 0.4483 | 29.67 | 0.7912 | 0.4649 |
| HMEC | chr20 | DiCARN | 32.78 | 0.8684 | 0.5214 | 34.56 | 0.8913 | 0.5258 |
| HMEC | chr20 | HiCARN1 | 34.53 | 0.8867 | 0.5070 | 34.65 | 0.8758 | 0.5859 |
| HMEC | chr20 | HiCNN | 27.06 | 0.8656 | 0.5051 | 28.39 | 0.8844 | 0.5070 |
| HMEC | chr20 | HiCPlus | 26.45 | 0.7629 | 0.4485 | 29.94 | 0.7942 | 0.4650 |
| *K562 (cross-cell, 1/16 LR)* | | | | | | | | |
| K562 | chr4 | DiCARN | 33.22 | 0.9412 | 0.8256 | 33.87 | 0.9440 | 0.8387 |
| K562 | chr4 | HiCARN1 | 33.11 | 0.9281 | 0.8048 | 34.02 | 0.9410 | 0.8254 |
| K562 | chr4 | HiCNN | 30.89 | 0.8498 | 0.7892 | 31.22 | 0.8550 | 0.6998 |
| K562 | chr4 | HiCPlus | 31.08 | 0.8198 | 0.7565 | 31.33 | 0.8585 | 0.7640 |
| K562 | chr14 | DiCARN | 33.65 | 0.9438 | 0.8301 | 34.23 | 0.9465 | 0.8423 |
| K562 | chr14 | HiCARN1 | 33.54 | 0.9307 | 0.8095 | 34.44 | 0.9435 | 0.8301 |
| K562 | chr14 | HiCNN | 31.23 | 0.8523 | 0.7943 | 31.54 | 0.8576 | 0.7047 |
| K562 | chr14 | HiCPlus | 31.42 | 0.8224 | 0.7612 | 31.66 | 0.8611 | 0.7688 |
| K562 | chr16 | DiCARN | 34.21 | 0.9465 | 0.8338 | 34.89 | 0.9492 | 0.8465 |
| K562 | chr16 | HiCARN1 | 34.08 | 0.9335 | 0.8132 | 35.01 | 0.9462 | 0.8336 |
| K562 | chr16 | HiCNN | 31.67 | 0.8549 | 0.7984 | 31.99 | 0.8601 | 0.7081 |
| K562 | chr16 | HiCPlus | 31.84 | 0.8248 | 0.7658 | 32.07 | 0.8637 | 0.7729 |
| K562 | chr20 | DiCARN | 34.48 | 0.9485 | 0.8381 | 35.13 | 0.9511 | 0.8521 |
| K562 | chr20 | HiCARN1 | 34.35 | 0.9353 | 0.8165 | 35.25 | 0.9487 | 0.8373 |
| K562 | chr20 | HiCNN | 31.81 | 0.8570 | 0.8013 | 32.17 | 0.8625 | 0.7112 |
| K562 | chr20 | HiCPlus | 31.94 | 0.8274 | 0.7673 | 32.18 | 0.8660 | 0.7751 |
| *NHEK (cross-cell, 1/16 LR)* | | | | | | | | |
| NHEK | chr4 | DiCARN | 34.78 | 0.9611 | 0.8431 | 35.02 | 0.9632 | 0.8485 |
| NHEK | chr4 | HiCARN1 | 34.72 | 0.9518 | 0.8146 | 35.01 | 0.9657 | 0.8101 |
| NHEK | chr4 | HiCNN | 33.68 | 0.8805 | 0.8116 | 34.67 | 0.9066 | 0.8116 |
| NHEK | chr4 | HiCPlus | 32.71 | 0.8468 | 0.8322 | 33.57 | 0.8665 | 0.7925 |
| NHEK | chr14 | DiCARN | 35.11 | 0.9628 | 0.8478 | 35.33 | 0.9648 | 0.8532 |
| NHEK | chr14 | HiCARN1 | 35.07 | 0.9534 | 0.8194 | 35.31 | 0.9674 | 0.8150 |
| NHEK | chr14 | HiCNN | 33.92 | 0.8821 | 0.8165 | 34.91 | 0.9082 | 0.8165 |
| NHEK | chr14 | HiCPlus | 33.02 | 0.8484 | 0.8371 | 33.89 | 0.8682 | 0.7974 |
| NHEK | chr16 | DiCARN | 35.48 | 0.9646 | 0.8510 | 35.71 | 0.9666 | 0.8564 |
| NHEK | chr16 | HiCARN1 | 35.44 | 0.9550 | 0.8228 | 35.68 | 0.9692 | 0.8182 |
| NHEK | chr16 | HiCNN | 34.23 | 0.8837 | 0.8198 | 35.29 | 0.9098 | 0.8198 |
| NHEK | chr16 | HiCPlus | 33.28 | 0.8499 | 0.8402 | 34.15 | 0.8698 | 0.8002 |
| NHEK | chr20 | DiCARN | 35.95 | 0.9659 | 0.8553 | 36.26 | 0.9674 | 0.8607 |
| NHEK | chr20 | HiCARN1 | 35.89 | 0.9562 | 0.8264 | 36.20 | 0.9701 | 0.8219 |
| NHEK | chr20 | HiCNN | 34.69 | 0.8857 | 0.8233 | 35.65 | 0.9118 | 0.8233 |

*Continued on next page*

Table S4 — *continued*

| Cell line | Chr | Generator | Standalone | | | + HiCP2GAN | | |
|---|---|---|---|---|---|---|---|---|
| | | | PSNR | SSIM | GDISCO | PSNR | SSIM | GDISCO |
| NHEK | chr20 | HiCPlus | 33.59 | 0.8513 | 0.8441 | 34.47 | 0.8711 | 0.8047 |

#### Supplementary Table S4: 1/16 vs. 1/64 low-resolution depth (GM12878)

The 1/16 columns repeat the chromosome-pooled *test* metrics from the main manuscript (same-cell GM12878 evaluation on chromosomes 4, 14, 16, and 20). The 1/64 columns report the same metrics computed on the 1/64 corpus (test chromosomes). **Supplementary Table S6** provides the per-chromosome breakdown for 1/64.

**Table S5. Supplementary Table S5.** GM12878 contact-map metrics under 1/16 vs. 1/64 downsampling (test chromosomes 4, 14, 16, 20).

| Setting | 1/16 **LR (test)** | | | 1/64 **LR (test)** | | |
|---|---|---|---|---|---|---|
| | PSNR | SSIM | GDISCO | PSNR | SSIM | GDISCO |
| DiCARN standalone | 35.72 | 0.9142 | 0.9138 | 32.84 | 0.8754 | 0.8795 |
| DiCARN + HiCP2GAN | 36.02 | 0.9213 | 0.9196 | 33.35 | 0.8886 | 0.8756 |
| HiCARN1 standalone | 35.64 | 0.9138 | 0.9106 | 33.61 | 0.8900 | 0.8905 |
| HiCARN1 + HiCP2GAN | 35.93 | 0.9202 | 0.9140 | 33.25 | 0.8873 | 0.8796 |
| HiCNN standalone | 35.46 | 0.9085 | 0.9079 | 28.01 | 0.3822 | 0.8342 |
| HiCNN + HiCP2GAN | 35.84 | 0.9161 | 0.9147 | 28.31 | 0.3822 | 0.8418 |
| HiCPlus standalone | 35.32 | 0.9051 | 0.9044 | 26.63 | 0.5650 | 0.8324 |
| HiCPlus + HiCP2GAN | 35.70 | 0.9124 | 0.9103 | 29.11 | 0.5812 | 0.8292 |

#### References


Suhas SP Rao, Miriam H Huntley, Neva C Durand, Elena K Stamenova, Ivan D Bochkov, James T Robinson, Adrian L Sanborn, Ido Machol, Arina D Omer, Eric S Lander, et al. A 3d map of the human genome at kilobase resolution reveals principles of chromatin looping. *Cell*, 159(7):1665–1680, 2014.

Xiao Wang, Yuanyuan Zhang, Suhita Ray, Anupama Jha, Tangqi Fang, Shengqi Hang, Sergei Doulatov, William Stafford Noble, and Sheng Wang. A generalizable hi-c foundation model for chromatin architecture, single-cell and multi-omics analysis across species. *bioRxiv*, 2024.
